# A complex phenotypic assay of mammalian oocyte maturation identifies compounds that block meiotic progression for non-hormonal contraceptive discovery

**DOI:** 10.1101/2025.10.15.682568

**Authors:** Jeffrey Pea, Yiru Zhu, Hoi Chang Lee, Lauren R. Haky, John Proudfoot, Richard Nelson, Francesca E. Duncan

## Abstract

Despite the prevalent use and efficacy of hormonal contraception, it remains unsuitable for many due to the prevalence of perceived and actual side effects. Thus, there is a need for novel contraceptives for women worldwide, especially non-hormonal options. During ovulation, oocytes undergo a maturation process by which they progress through meiosis, a specialized cell division needed to reduce chromosome content. Given that meiotic progression is essential for generating a fertilizable egg but does not affect hormone production, it represents a promising biological mechanism to target for female non-hormonal contraception. In this study, we developed a phenotypic screening platform modeled after biopharmaceutical industry practices using mouse oocytes to screen for compounds that block meiotic progression. Oocytes were matured *in vitro* in the presence of selected compounds from a bioactive compound library, and confirmed hits were identified as those that potently and reproducibly blocked meiotic progression. In addition, we showed that the inhibitory effect of these compounds was persistent and reversible. Thus, our oocyte drug screening pipeline provides a powerful system to identify novel drug discovery candidates for further follow up for non-hormonal contraception development.

## Introduction

Modern methods of contraception are an essential component of reproductive and sexual health care with their use doubling up to 874 million women worldwide in the past three decades (United Nations Department of Economic and Social Affairs, Population Division, 2022). Despite this increased prevalence, there is a gap in access to contraceptives with approximately 121 million unintended pregnancies occurring annually due in part to unmet family planning needs (Sully *et al*., 2020). For many women, the current options for contraception, especially hormonal-based methods, are not suitable due to associated side effects, health risks, and inaccessibility (Sedgh *et al*., 2016). In fact, despite high efficacy, hormonal contraception has significant side effects, including risk of venous thrombotic events, migraines with aura, fatigue, and mood swings (Teal and Edelman, 2021), with up to 40% of women in low and middle income countries discontinuing use within the first year (Safari *et al*., 2019; Rothschild *et al*., 2022). As such, there is a clear need to develop additional contraceptive options, in particular non-hormonal contraception.

In the ovary, in response to the surge of luteinizing hormone that triggers ovulation, oocytes within the ovulatory follicle undergo a final maturation process that is essential to generate a fertilizable gamete. During this process, oocytes, which are arrested in prophase of meiosis I throughout the reproductive lifespan, resume and complete meiosis I. Oocytes then arrest again at metaphase of meiosis II (MII) until fertilization triggers completion of meiosis II. Notably, only eggs that are arrested at MII can be fertilized by sperm, and inhibition of meiotic progression does not impact estrous cyclicity, ovulation, or hormone production (Masciarelli *et al*., 2004a). Furthermore, meiotic progression is an oocyte-specific process that only occurs within a very narrow timeframe of ovulation. As such, blocking meiotic progression offers a promising biological mechanism-of-action for non-hormonal contraception.

Conventional drug discovery pipelines fall into two primary approaches: target-based and phenotypic-based. Whereas target-based drug discovery starts with a defined molecular target and has recently been the dominant approach in the pharmaceutical industry, phenotypic drug discovery is target-agnostic and can identify active molecules that modulate a specific process within a physiologically relevant biological system (Moffat *et al*., 2017). In other fields such as infectious disease, phenotypic drug discovery strategies (e.g., viral / bacterial replication inhibition) have proven successful in translation into *in vivo* preclinical and clinical therapies (Andries *et al*., 2005; Ma *et al*., 2007). However, there remains an absence of robust cell-based phenotypic assays within the field of reproductive biology with few being leveraged for screening potential contraceptive drug candidates (Gruber *et al*., 2020; Zhang *et al*., 2023).

Oocyte maturation is an ideal biological process to adapt to drug screening because phenotypic changes can be fully captured within a cell-based system. Notably, mouse oocytes undergo spontaneous meiotic maturation *in vitro* once they are removed from the follicle (Pincus and Enzmann, 1935). In addition, meiotic stages can be assessed by morphological criteria that are readily visible via transmitted light microscopy (Suebthawinkul *et al*., 2022, 2023; Zhu *et al*., 2024). For example, the presence of the oocyte nucleus or germinal vesicle (GV) is indicative of prophase of meiosis I, the presence of the first polar body (PB) is indicative of arrest at MII, and the absence of both the GV and PB is indicative of pro-metaphase or metaphase of meiosis I (Suebthawinkul *et al*., 2022, 2023; Zhu *et al*., 2024). These features provide a unique opportunity to harness mouse oocyte maturation as a complex phenotypic assay that can be used to screen drugs that block meiotic progression for contraceptive discovery purposes.

Therefore, in this study we established and validated an oocyte maturation assay, and then screened 818 compounds selected from a structurally diverse and medicinally active library of biologically active molecules to identify those that blocked meiotic progression. Initial hits were subject to additional validation approaches, including confirmation with an independent compound source and establishment of a concentration-dependent response. Hit compounds were further evaluated in a counterscreen for PDE3A activity, for their ability to maintain prolonged meiotic arrest, and in reversibility studies of pharmacological inhibition. Finally, initial structure activity relationships were probed through comparisons of structural analogs. An overview schematic summarizing this phenotypic screening pipeline is presented in **Fig. 1**.

**Figure 1.**
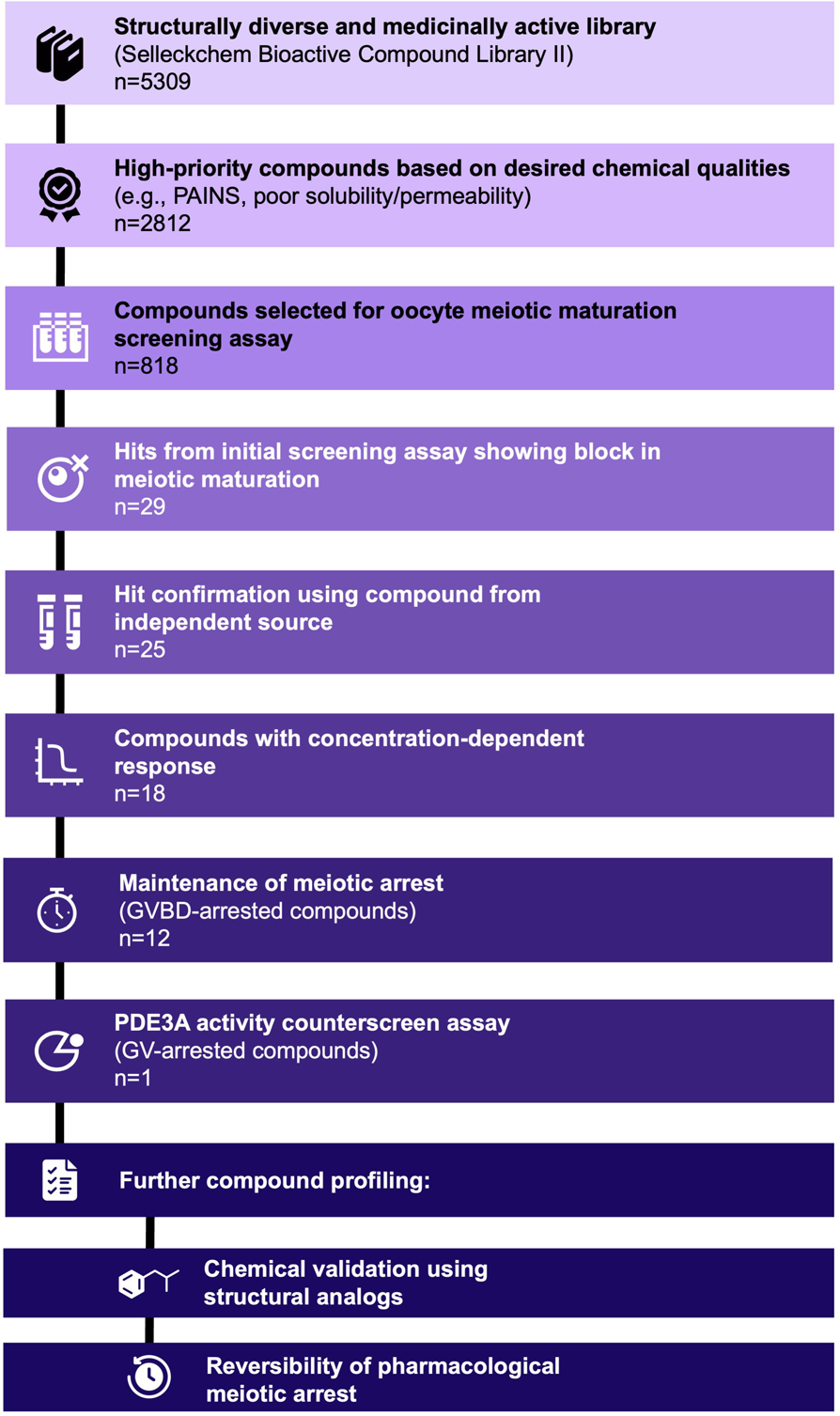
Overview of compounds screened within murine oocyte meiotic maturation assay and screening pipeline. Compounds from a bioactives library (Selleckchem Bioactives Library II) were filtered to remove those with undesirable chemical properties and prioritize a subset for testing within the phenotypic screening assay. Hits identified from the initial counterscreen assay then underwent hit validation, including confirmation using an independent source, concentration-response, and prolonged culture treatment. For GV-arresting compounds, an additional PDE3A activity counterscreen assay was conducted as well as further compound profiling, including chemical validation using structural analogs and reversibility studies.

## Materials and Methods

### Chemical and library selection

A library with 5,309 medicinally active, structurally diverse, and cell permeable compounds was purchased from Selleckchem (Bioactives Library II). However, compounds were filtered to remove those flagged with pan-assay interference (PAINS) activity, therefore reducing assay specificity and generating false positive assay results, or based on chemical features that are conventionally unprogressible in drug development (BAD-SMARTS). After filtering out compounds with undesirable qualities, including PAINS activities, BAD-SMARTS features, chemically reactive features, calculated cell permeability, and poor solubility and permeability, we obtained a list of 2,812 high-priority compounds (**Fig. 1**, **Fig. 2A, Supplemental Fig. 1**). From this subset, we deprioritized out compounds annotated to bind to sex hormone (androgen, estrogen) receptors or affect sex hormone (androgen, estrogen, progesterone) production and selected a final set of 818 compounds based on annotated target and chemical structural diversity (**Fig. 2B**). These compounds were supplied in “assay-ready” plates and solubilized prior to their addition to culture medium. All the compounds were pre-dissolved in DMSO as a 10 mM stock solution and divided into 5 μl aliquots stored at −80 °C. The primary screening assay used compound stock aliquots stored at room temperature for no longer than 3 months after thawing.

**Figure 2.**
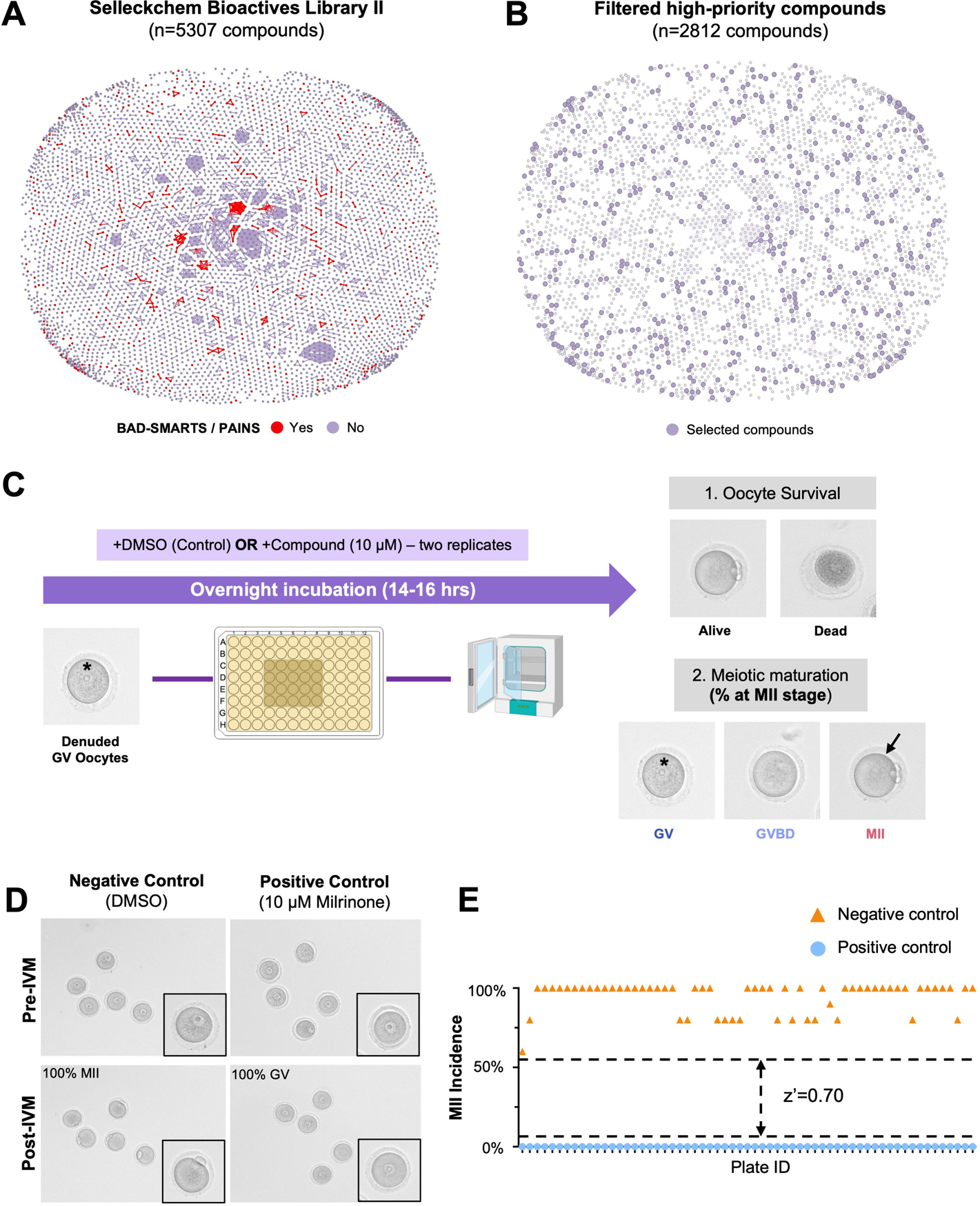
Compound selection and murine oocyte meiotic maturation assay. (**A**) Structural similarity chart of compounds from the entire Selleckchem Bioactives Library II (n=5307). Red dots indicate compounds deprioritized due to unprogressable (BAD-SMARTS) or pan-assay interference (PAINS) features. (**B**) Structural similarity chart of high-priority compounds filtered from the Selleckchem Bioactives Library (n=2812). Purple dots indicate the final compounds selected for the phenotypic oocyte maturation screening assay. (**C**) Overview of phenotypic oocyte maturation screening assay. Denuded oocytes at the GV stage were placed in an overnight incubator for in vitro maturation (IVM) with either DMSO (negative control), 10 µM Milrinone (positive control), or 10 µM compound-of-interest. Brightfield images were taken before and after IVM to determine oocyte survival and maturation status. Asterisk indicates germinal vesicle and black arrow indicates the extruded polar body. (**D**) Representative images of oocytes treated with DMSO (negative control) and 10 µM Milrinone (positive control) before and after IVM. (**E**) Graph of MII incidence for negative and positive controls across phenotypic oocyte screening assay plates and of Z’-factor to determine assay quality. GV, germinal vesicle; GVBD, germinal vesicle breakdown; MII, meiosis II.

### Animals

Reproductively adult CD1 female mice (6-12 weeks of age) were obtained from Envigo (Indianapolis, IN). They were housed in a controlled barrier facility at the Northwestern University Center for Comparative Medicine under constant temperature, humidity, and light (14 h light/10 h dark). Mice were provided with water and Teklad Global 2916 chow (Envigo) *ad libitum*, a diet that excludes soybean meal. The mice were allowed to acclimate for at least a week prior to use in experiments. All animal experiments described were approved by the Institutional Animal Care and Use Committee (Northwestern University) and performed under the National Institutes of Health Guidelines.

### Ovarian hyperstimulation and oocyte collection

Mice were administered intraperitoneal injections of 5 IU pregnant mare serum gonadotropin (PMSG) (ProSpec-Tany TechnoGene, East Brunswick, NJ, Cat # HOR-272) 44– 46 hours prior to harvest to optimize oocyte yield. Ovaries were dissected and placed into dishes with pre-warmed Leibovitz’s medium (L15) (Life Technologies Corporation, Grand Island, NY) supplemented with 3 mg/ml polyvinylpyrrolidone (PVP) (Sigma-Aldrich, St. Louis, MO) and 0.5% (v/v) of Penicillin–Streptomycin (PS) (Life Technologies Corporation, Grand Island, NY) (L15/PVP/PS). Cumulus-oocyte-complexes (COC) were obtained by mechanically puncturing the antral follicles with a syringe needle. The COCs were subsequently moved to L15/PVP/PS media supplemented with 2.5 μM milrinone (Sigma-Aldrich, St. Louis, MO), an inhibitor of phosphodiesterase 3A (PDE3A) to maintain oocyte arrest at prophase of meiosis I (Wiersma *et al*., 1998). To obtain denuded oocytes, the cumulus layer of COCs was removed by mechanical disruption with a stripper tip. All the GV oocytes were then allowed to recover in a dish with pre-equilibrated α-MEM+GlutaMAX (Thermo Fisher Scientific, Waltham, MA, USA)/0.5% PS /0.3% bovine serum albumin (BSA) (Sigma-Aldrich) (α-MEM/PS/BSA) media supplemented with 2.5LμM milrinone for 1 h at 37°C in a humidified atmosphere of 5% CO_2_ in air to recover from the isolation process while maintaining meiotic arrest prior to experimental use. Oocytes from eight mice per experiment were pooled together to minimize any animal-specific variability.

### Oocyte maturation screening assay

For the primary screening assay, 22 wells of the 96-well flat-bottom microplates (Corning; 3598) were loaded with 100 μl α-MEM/BSA/PS media containing 10 μM compound, 10 μM milrinone (positive control), or dimethyl sulfoxide (DMSO) (Sigma-Aldrich, St. Louis, MO, Cat # M1404) of equal volume as the vehicle control in the absence of an oil overlay (**Fig. 2C**). The starting concentration of 10µM was chosen as it is considered standard practice in single-dose primary drug screen pipelines (Hughes *et al*., 2011). The rest of the wells were filled with sterile Dulbecco’s phosphate-buffered saline (DPBS) without calcium and magnesium (Thermo Fisher Scientific, Waltham, MA) to maintain humidity. The microplates were allowed to equilibrate for at least 4 hours in the incubator prior to screening. To initiate oocyte maturation, denuded GV oocytes were washed in 4-well dishes (Thermo Fisher Scientific, Waltham, MA) containing pre-equilibrated α-MEM/BSA/PS media without milrinone. The removal of milrinone results in the degradation of cyclic adenosine monophosphate (cAMP) and spontaneous meiotic resumption. Five oocytes were loaded into each treatment well of the 96-well microplate and IVM was performed for 14-15 h at 37 °C in a humidified atmosphere of 5% CO2 in air. Images were taken before and after incubation using the EVOS FL Auto Imaging System with a 10X objective (Thermo Fisher Scientific) to examine oocyte morphology and maturation status. The washing and loading of oocytes were conducted at the same time by a single operator. Following IVM, the maturation status of oocytes was determined based on established morphological features (Suebthawinkul *et al*., 2022, 2023; Zhu *et al*., 2024). Oocytes that underwent maturation and arrested at the MII stage were characterized by the extrusion of the first PB. Oocytes that had an intact GV were arrested in prophase of meiosis I and referred to as GV oocytes. Oocytes that progressed through meiosis and extruded a polar body were considered at the MII stage. Oocytes that lacked a GV and had undergone germinal vesicle breakdown (GVBD) but lacked a PB were considered in pro-metaphase I or metaphase I and thusly referred to as GVBD oocytes. Following IVM, the presence of GV oocytes and GVBD oocytes were indicative of blocked meiotic progression. Oocytes that had a darkened or flat appearance were considered degenerate as a result of cytotoxicity (**Fig. 2C**). All compounds were screened in duplicate, and the percentage of oocytes at each meiotic stage was calculated per trial. A hit within the primary screening assay was defined as a compound that blocked meiotic progression by at least 80% in both trials.

### Concentration response and prolonged culture treatment

Following identification of hits from the primary screening assay, oocytes were treated with a 3-fold serial dilution of each hit compound, starting from 10 μM as the highest concentration, to determine if there is a concentration-dependent effect. A minimum of five concentrations were evaluated for each compound in the concentration-response analysis. Following confirmation of concentration-dependent response, oocytes were also treated at 10 µM of each compound for an extended IVM time course of up to 38 h. Images were taken before and during the incubation period at 14 h (conventional IVM endpoint), 26 h, and 38 h. For these experiments, DMSO and 10 µM milrinone were used as negative and positive controls, respectively.

### PDE3A activity assay

PDE3A has previously been identified as a promising non-hormonal contraceptive target whose inhibition blocks oocyte maturation without affecting ovulation or estrous cyclicity (Wiersma *et al*., 1998; Li *et al*., 2012). As such, a PDE3A activity assay was conducted as a counterscreen assay in an effort to focus on novel targets. Specificially, PDE3A enzymatic activity was determined using a human PDE3A AMP-Glo^TM^ assay (Pharmaron, Beijing, China) which detects AMP levels via luciferase reaction following enzymatic cleavage of cAMP to AMP. Compounds that maintained GV arrest were tested at 10 concentrations at 3-fold dilutions from 10 µM to 0.51 nM. Cilostamide, a potent PDE3 inhibitor, was used as the reference compound for the assay. IC50 values were then calculated to determine the concentration-dependent effect of the compounds on PDE3A activity.

### Pharmacologic reversibility analysis

One major factor to consider for non-hormonal contraceptive drug development is reversibility and ensuring that oocytes remain healthy and viable following drug discontinuation. As such, a reversibility study was conducted to determine oocyte maturation status before and after compound washout. For the first incubation, oocytes were treated with 10 µM compound within a 96-well microplate and IVM was performed for 14-15 h as previously described for the initial primary screen. Following IVM, treated oocytes were washed four times in 800 µl of L15/PVP/PS media to remove the compound. In order to determine whether or not oocyte maturation progresses normally following compound washout, a closed timelapse incubator (EmbryoScope+ ^TM^) was used evaluate timing and morphokinetic parameters of meiotic progression as previously described (Suebthawinkul *et al*., 2022, 2023; Zhu *et al*., 2024).

Following washout, oocytes were placed into the EmbryoSlides with milrinone-free media and incubated the EmbryoScope+^TM^ system for 14-15 h. EmbryoSlides were prepared the day before the experiment following manufacturer’s instructions. Each microwell was filled with α-MEM/BSA/PS media and overlayed with 1.6 ml of mineral oil (Sigma-Aldrich, St. Louis, MO) before equilibration for 9–11 h. Images were taken every 10 minutes to monitor the maturation status of the oocytes. Morphokinetic parameters of meiotic progression, including time to GVBD, time to PBE, and duration of MI, were determined as previously described (Suebthawinkul *et al*., 2022) for all oocytes arrested at both GV and GVBD stages. In brief, t0 was defined as the time when denuded oocytes were placed into the EmbryoScope+^TM^ and imaging was started. Time to GVBD was defined as the time when the complete loss of the first GV was first observed and time to PBE was defined as the time when there was clear separation between the polar body and oocyte membranes. Duration of MI was defined as the time difference between time to GVBD and time to PBE. Oocytes treated with 10 µM milrinone in the first incubation but washed and moved into milrinone-free media in the second incubation served as positive controls.

### Immunofluorescence and microscopy

Following the prolonged culture treatment and reversibility studies, cytoskeletal morphology was evaluated in oocytes using immunofluorescence and microscopy. Oocytes were fixed in 3.8% paraformaldehyde (Electron Microscopy Sciences, Hatfield, PA, USA) with 0.1% Triton X-100 (TX-100) (Sigma-Aldrich) for 25 min at 37 °C followed by washing with blocking buffer (1x PBS, 0.01% Tween-20 (Sigma-Aldrich), 0.02% sodium azide (NaN_3_) (Sigma-Aldrich), and 0.3% BSA) four times (5 min each wash). They were then transferred to permeabilization solution (1x PBS, 0.1% TX-100, and 0.3% BSA) for 15 min at room temperature. Oocytes were washed again with blocking buffer twice (5 min each wash) and subsequently incubated with rhodamine phalloidin (1:50; Invitrogen, Waltham, MA, USA, Cat No. R415) and α-tubulin (11H10) Rabbit mAB 488 (1:100; Cell Signaling Technology, Danvers, MA, USA, Cat No. 5063S) for 2 h at room temperature to visualize actin and microtubules, respectively. Afterwards, oocytes were rinsed with blocking buffer three times (20Lmin each wash) and mounted on slides in Vectashield Antifade Mounting Medium with DAPI (4, 6-diamidino-2-phenylindole; Vector Laboratories, Burlingame, CA, USA). Cells were imaged at 40× magnification on a Leica SP5 inverted laser scanning confocal microscope (Leica Microsystems, Wetzlar, Germany) using 405, 488, and 543 nm lasers. All images were processed using LAS AF (Leica Microsystems) and analyzed using FIJI (National Institutes of Health, Bethesda, MD, USA).

### Statistical analysis

Each experiment was repeated at a minimum of two times. Results were generated using GraphPad Prism Software Version 8.0.1 (La Jolla, CA, USA). Z-prime factor (Z’-factor) is a parameter commonly used during high-throughput screening to determine the quality of its assays (Zhang *et al*., 1999). Z’-factor measures statistical effect size by comparing the response (means and standard deviations) of the positive and negative controls and is scaled from 0.0 to 1.0. Assays with a Z’-factor >0.5 are considered “excellent” and sufficient for high-throughput screening. IC50 represents the concentration of compound that is required to inhibit a biological process (e.g., MII incidence) by 50%. IC50 values and concentration response curves were generated using a non-linear regression model with a minimum of five concentrations with the fixed bottom parameter (lower asymptote) = 0%. The normal distribution of data was evaluated with the Shapiro–Wilk test. Analysis between groups of continuous variables was performed with Student’s *t*-test.

## Results

### Phenotypic oocyte maturation assay and primary screen

A primary cell-based assay was optimized in a 96-well plate format to allow for IVM within a conventional incubator and without an oil overlay to identify compounds that block meiotic progression without affecting oocyte viability (**Fig. 2C**). Through this approach, we identified phenotypes across meiotic progression, including maintained arrest at the germinal vesicle stage (GV), germinal vesicle breakdown but failure to reach the MII arrest (GVBD), and progression to the MII stage (MII) (**Fig. 2C**). Degenerate cells were also distinguished based on morphology (**Fig. 2C**). As part of the assay, we established baseline parameters with both negative (DMSO, vehicle) and positive controls (Milrinone, a PDE3 inhibitor). As expected, untreated oocytes progressed to the MII stage whereas oocytes treated with Milrinone exhibited arrest at the GV stage (**Fig. 2D**). As part of assay development for high-throughput screening, standard quality control procedures include validating that the assay can robustly differentiate between the positive and negative controls. Indeed, the oocyte maturation assay had a Z’-factor = 0.70, indicating high assay quality (**Fig. 2E**). We also tested this system using a series of known cytotoxic or pan-assay interfering (PAINS) compounds to determine their ability to cause oocyte toxicity and non-selective hits, respectively (**Supplemental Fig. 2**). PAINS are undesirable chemical compounds that non-selectively react in various biological assays and must be considered in drug discovery pipelines (Baell and Holloway, 2010; Chakravorty *et al*., 2018). Indeed, oocyte maturation in the presence of known PAINS compounds resulted in undesired phenotypes, such as degeneration, as well as meiotic arrest phenotypes (**Supplemental Fig. 2A, 2B**). Thus, in the Selleckchem Bioactives Compound Library II used in this study, we filtered and removed PAINS and cytotoxic compounds to prevent detection of undesired, non-selective hits during screening (**Fig. 2A**).

Ultimately after this filtering, a subset of 818 compounds was further selected based on their diverse set of annotated targets and compound structures (**Fig. 2B**). Hits are defined within the phenotypic screening assay if the compounds blocked meiotic progression by at least 80% across two trials (**Fig. 3A, 3B**). Compounds were deprioritized if they exhibited oocyte degeneration or abnormal oocyte morphology due to toxicity or potential function as a PAINS compound. From our initial hits, we further prioritized compounds that maintained meiotic arrest at the GV stage and deprioritized those that underwent GVBD but maintained MI arrest (GVBD) (**Fig. 3A, 3B)**. In total, 29 hits were identified from the primary screen for an initial hit rate of 3.55% (**Fig. 1**, **Fig. 3C**).

**Figure 3.**
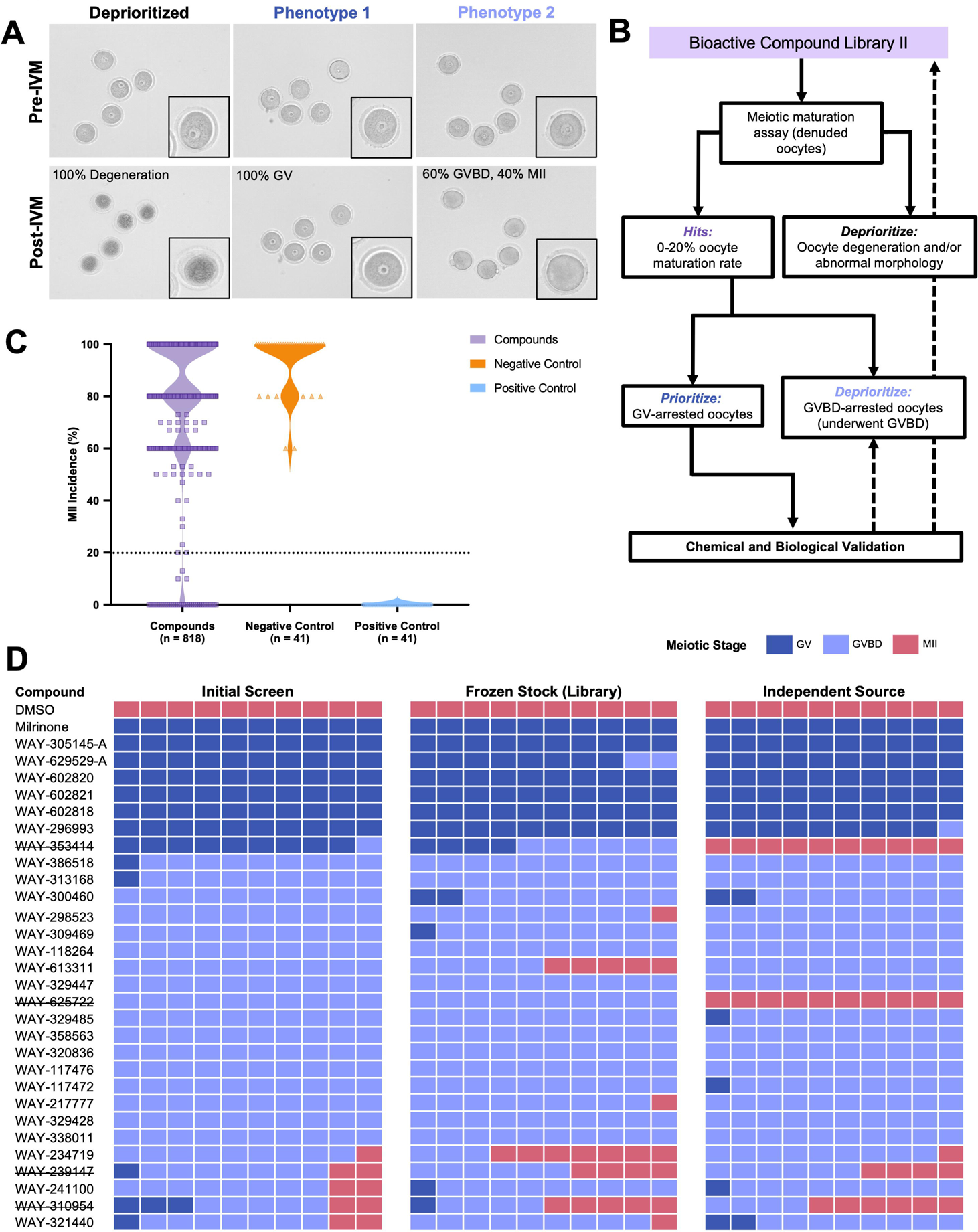
Results from initial phenotypic oocyte maturation screening assay and subsequent hit validation using independent source. (**A**) Representative images of oocyte phenotypes observed within initial screening assay, including deprioritized (oocyte degeneration), GV arrest, and GVBD arrest. (**B**) Decision tree for hit definition within oocyte maturation assay and hit prioritization for chemical and biological validation. (**C**) Graph of MII incidence of all tested compounds from initial oocyte maturation assay. Dotted line indicates cutoff for hit definition (≤20%). (**D**) Comparison of oocyte maturation status across treatment conditions, including initial library stock, frozen library stock, and source from independent vendor. Each box represents one oocyte. Crossed out compounds indicate those whose results from independent source did not align with results from library stocks. GV, germinal vesicle; GVBD, germinal vesicle breakdown; MII, meiosis II.

### Primary screen validation

Next, we validated the initial hits using two standard approaches in the high-throughput drug screening industry: (1) confirmation of findings with an independent source of compound and (2) concentration responsiveness. Of the total 29 compounds that initially blocked meiotic progression, 25 of them recapitulated results using an independent compound source and were analyzed further (**Fig. 3D**). These compounds were then tested using a standardized concentration range of 0.1, 0.3, 1, 3, and 10 µM to determine concentration-dependent effects on meiotic progression. For potent compounds that primarily blocked meiotic progression at 0.1 and 0.3 µM, lower concentrations were also tested to calculate IC50 values. A total of 18 out of 25 compounds blocked meiotic progression in a concentration-dependent manner (**Fig. 4A**). The remaining 7 compounds only elicited an effect at 10 µM and thus were likely non-specific and deprioritized (**Fig. 4B**).

**Figure 4.**
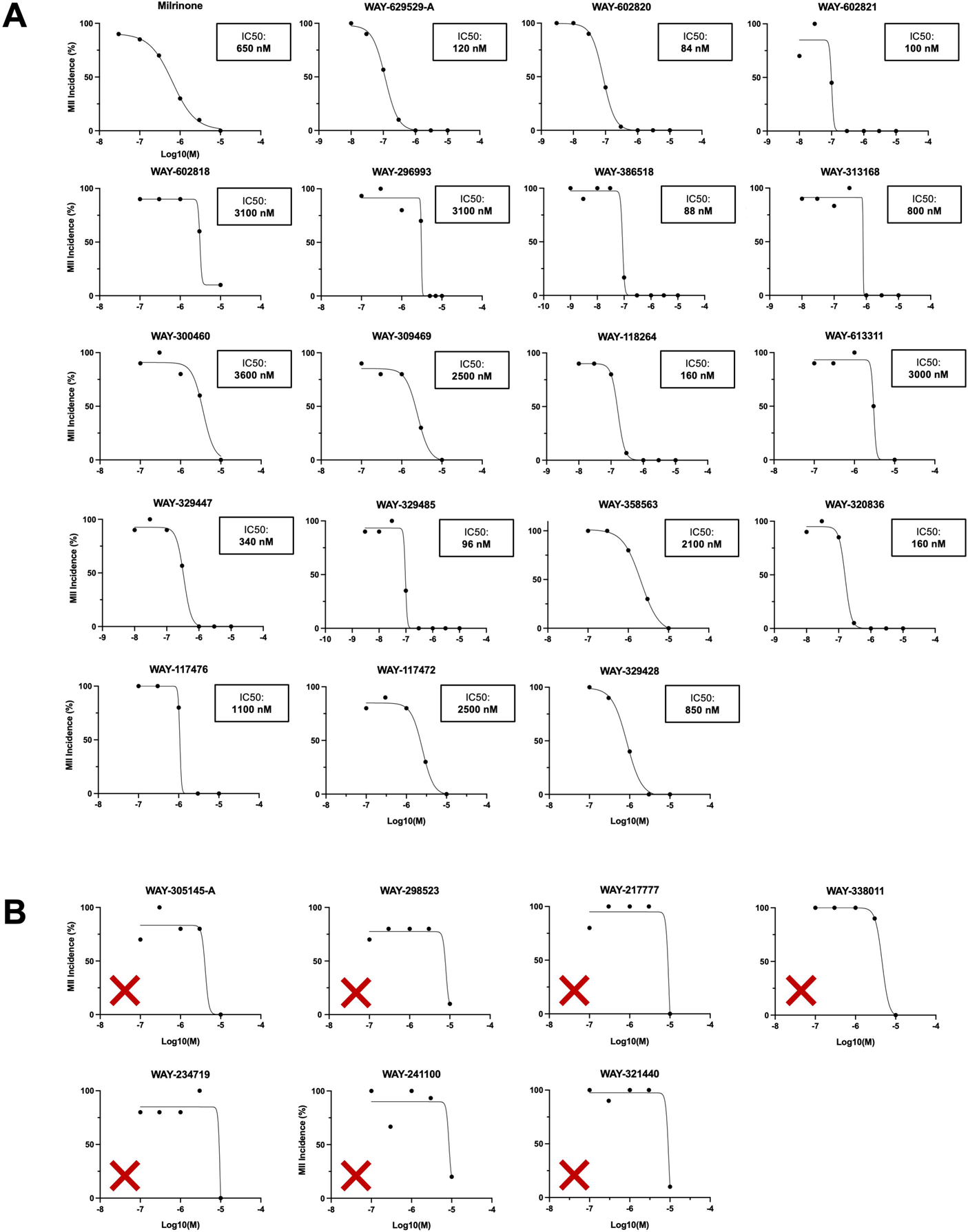
Hit validation via concentration-dependent response. (**A**) Concentration response curves and calculated IC50 for MII incidence in response to hit compounds. (**B**) Concentration response curves for MII incidence for hit compounds that did not exhibit concentration-dependent effect). MII, meiosis II.

Of the resulting 18 compounds, 13 resulted in GVBD arrest. Following ovulation, the mature egg typically remains viable and can be fertilized up to 24 h after reaching the MII arrest. Thus, we wanted to confirm that cells treated with these compounds do not eventually reach the MII stage beyond the window of IVM (14-15 h) as this could lead to breakthrough fertilization in an *in vivo* setting. To do so, we cultured oocytes for a prolonged period of up to 38 hours in the presence of the 13 compounds that resulted in GVBD arrest. The meiotic maturation status of oocytes was monitored at 0 h, 14 h, 26 h, and 38 h. The majority of the compounds (12 of 13) maintained meiotic arrest, with only one compound reaching the MII stage by 26 h (**Fig. 5A**). Following the prolonged culture, we fixed and stained the cells to evaluate chromosome configuration and cytoskeleton morphology (**Fig. 5B**). Most compounds (9/13) exhibited scattered chromosomes, enrichment of actin surrounding DNA at the cortex, and absence of tubulin suggesting the inability to generate and maintain a normal bipolar meiotic spindle with chromosomes aligned on the metaphase plate (**Fig. 5B**). However, one compound exhibited mini-spindles formed around each cluster of DNA (WAY-117472) and another compound exhibited tightly condensed chromosomes but scattered actin foci throughout the cytoplasm (WAY-613311) (**Fig. 5B**). Although cells cultured in WAY-300460 were able to reach the MII stage after extended culture, the meiotic spindles were abnormal, lacking a bipolar spindle and aligned chromosomes. These results suggest that compounds that arrest oocytes in the GVBD state likely do so through modulation of the oocyte cytoskeleton and prevention of proper meiotic spindle assembly or maintenance.

**Figure 5.**
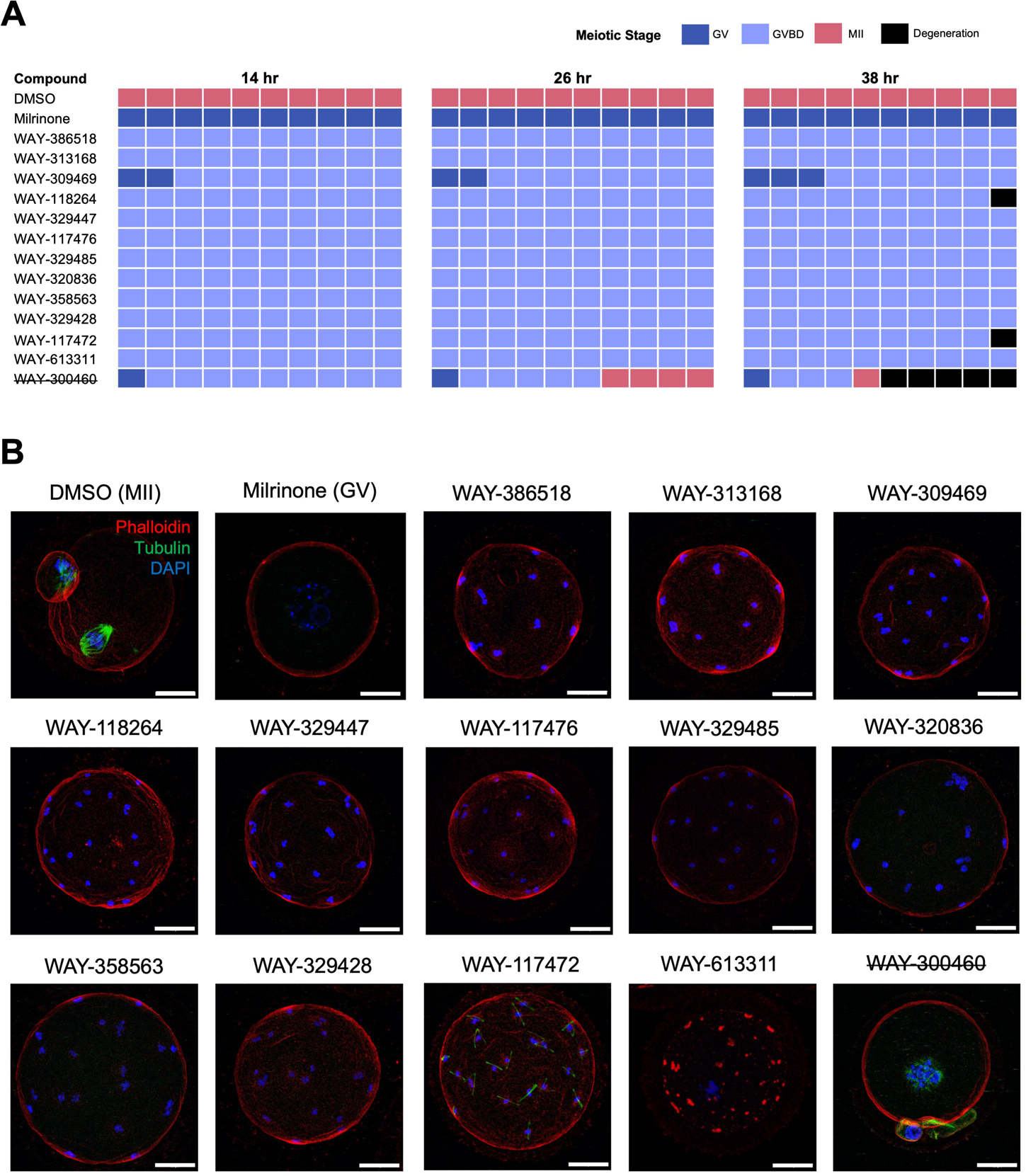
Hit validation via prolonged culture treatment. (**A**) Comparison of oocyte maturation status for each hit compound at regular intervals (14, 26, 38 hr) throughout prolonged culture treatment. Each box represents one oocyte. Crossed out compounds indicate those that did not maintain meiotic arrest throughout treatment window. (**B**) Representative immunofluorescent images showcasing the spindle morphology of compound-treated oocytes at the end of 38 hr treatment period. Adjustments were made to highlight localization/distribution of structures within the cell. Scale bar = 25 µm. MII, meiosis II; GV, germinal vesicle.

In addition to identifying compounds that blocked meiotic progression at the GVBD stage, the screening pipeline also identified 5 compounds that resulted in GV arrest. Such compounds are ideal for contraceptive development since cells arrested at prophase of meiosis I, even if ovulated, are unable to be fertilize. However, a major known meiotic regulator is PDE3A which is enriched in the oocyte, and its inhibition keeps cyclic AMP levels elevated and maintains prophase I arrest (Wiersma *et al*., 1998; Li *et al*., 2012). In fact, mice that lack oocyte specific PDE3A are infertile due to ovulation of oocytes arrested at the GV stage which cannot be fertilized (Masciarelli *et al*., 2004b). To rule out our hit compounds that were functioning through PDE3A, the compounds were tested against PDE3A via an enzymatic counterscreen assay. Four of these 5 compounds, all of which shared the same chemical substructure, exhibited concentration-dependent activity for PDE3A and thus were deprioritized for further validation (**Fig. 6A**). As such, only one compound (WAY-296993) elicited a GV-arrest phenotype without affecting PDE3A activity. Although less potent than the other compounds that maintained GV arrest (IC50: 3080 nM), WAY-296993 exhibited a robust concentration response pattern (**Fig. 6B**). Additionally, prolonged treatment with WAY-296993 at 5 µM exhibited persistence in preventing oocyte maturation even after 38 h of culture (**Fig. 6C**).

**Figure 6.**
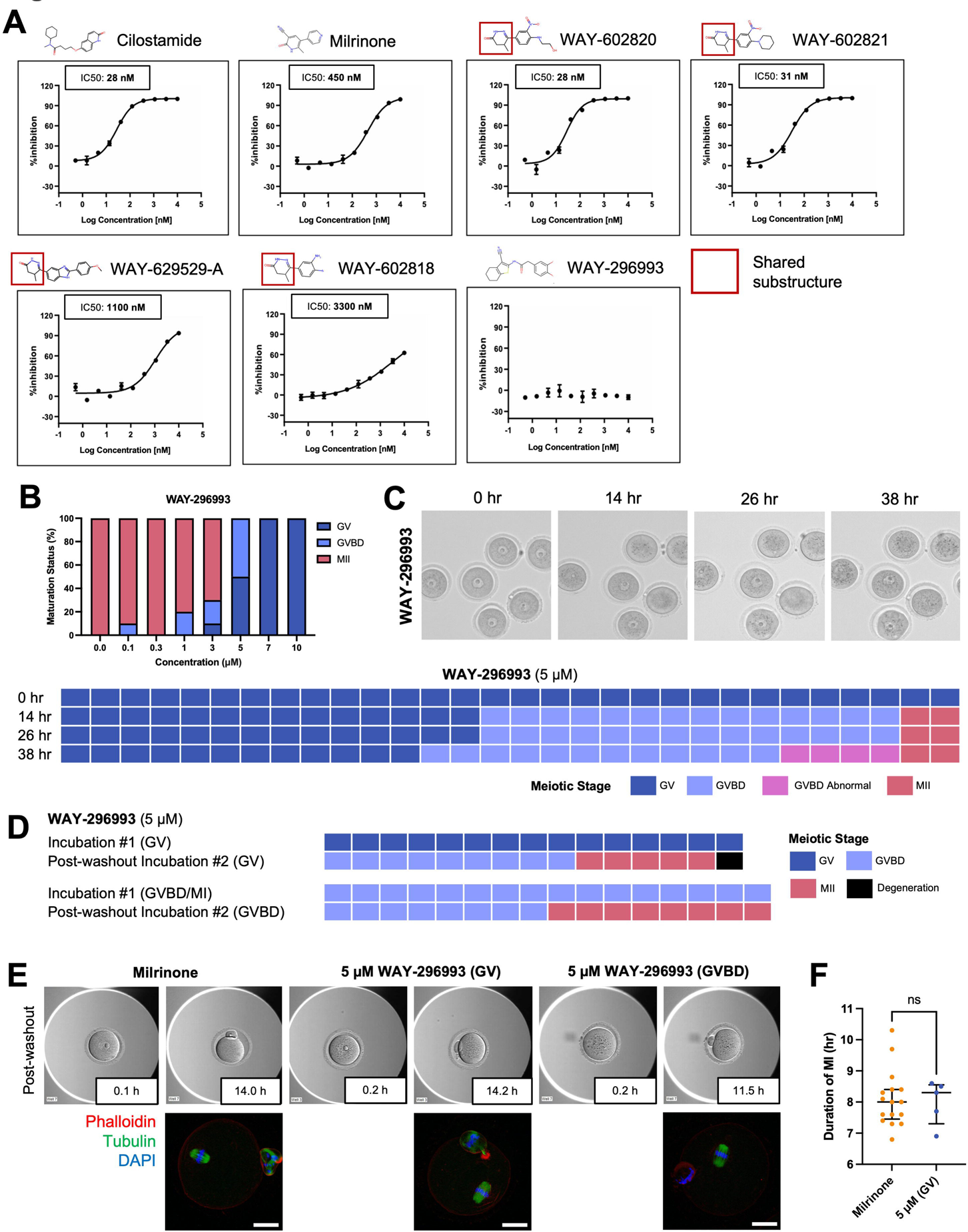
PDE3A activity counterscreen assay and additional compound profiling of GV-arresting hit compounds. (**A**) Concentration response curves for PDE3A activity in response to treatment with GV-arresting hit compounds. Red box denotes shared chemical substructure between compounds. (**B**) Graph of meiotic maturation status of WAY-296993 across different concentrations. Each box represents one oocyte. (**C**) Representative images and comparisons of oocytes treated with 5 µM WAY-296993 at regular intervals (14, 26, 38 hr) throughout prolonged culture treatment. (**D**) Comparison of oocyte maturation status between GV-arrested and GVBD-arrested oocytes treated with 5 µM WAY-296993. Each box represents one oocyte. (**E**) Representative EmbryoScope+^TM^ images of oocytes during second incubation following washout of either 10 µM Milrinone or 5 µM WAY-296993 treatment during first incubation period (top panel). Representative immunofluorescent images showcasing spindle morphology of oocytes from each treatment group at the end of the second incubation period (bottom panel). Scale bar = 25 µm. (F) Graph of duration of MI between GV-arrested oocytes treated with 5µM WAY-296993 and DMSO-treated oocytes. GV, germinal vesicle; GVBD, germinal vesicle; MI, meiosis I; MII, meiosis II.

An additional major consideration for compounds being developed for non-hormonal contraception is their reversibility. As such, we conducted a reversibility study with WAY-296993 in which oocytes were first treated with compound, rinsed to remove the compound, and subsequently cultured for another incubation period to track meiotic resumption. At 5 µM, WAY-296993 generated oocytes that were arrested both at the GV stage or GVBD stage following standard IVM (**Fig. 6B**). Following washout, a subset of oocytes of both phenotypes were able to resume meiosis during the second incubation and reach the MII stage (**Fig. 6D**). In oocytes that were able to progress to the MII stage, they showed a fully extruded polar body and normal spindle morphology (**Fig. 6E**). In addition, oocytes arrested at the GV stage by WAY-296993 exhibited no difference in time to PBE and duration of MI following drug washout relative to control oocytes arrested at the GV stage by Milrinone (**Fig 6.F**). These findings demonstrate that pharmacological inhibition of oocyte maturation can be reversible. In summary, our comprehensive compound screening pipeline identified 13 compounds that block meiotic progression with an overall hit rate of 1.6% (**Fig. 7**). We identified one compound that resulted in GV arrest and the remaining 12 resulting in GVBD arrest (**Table 1**).

**Figure 7.**
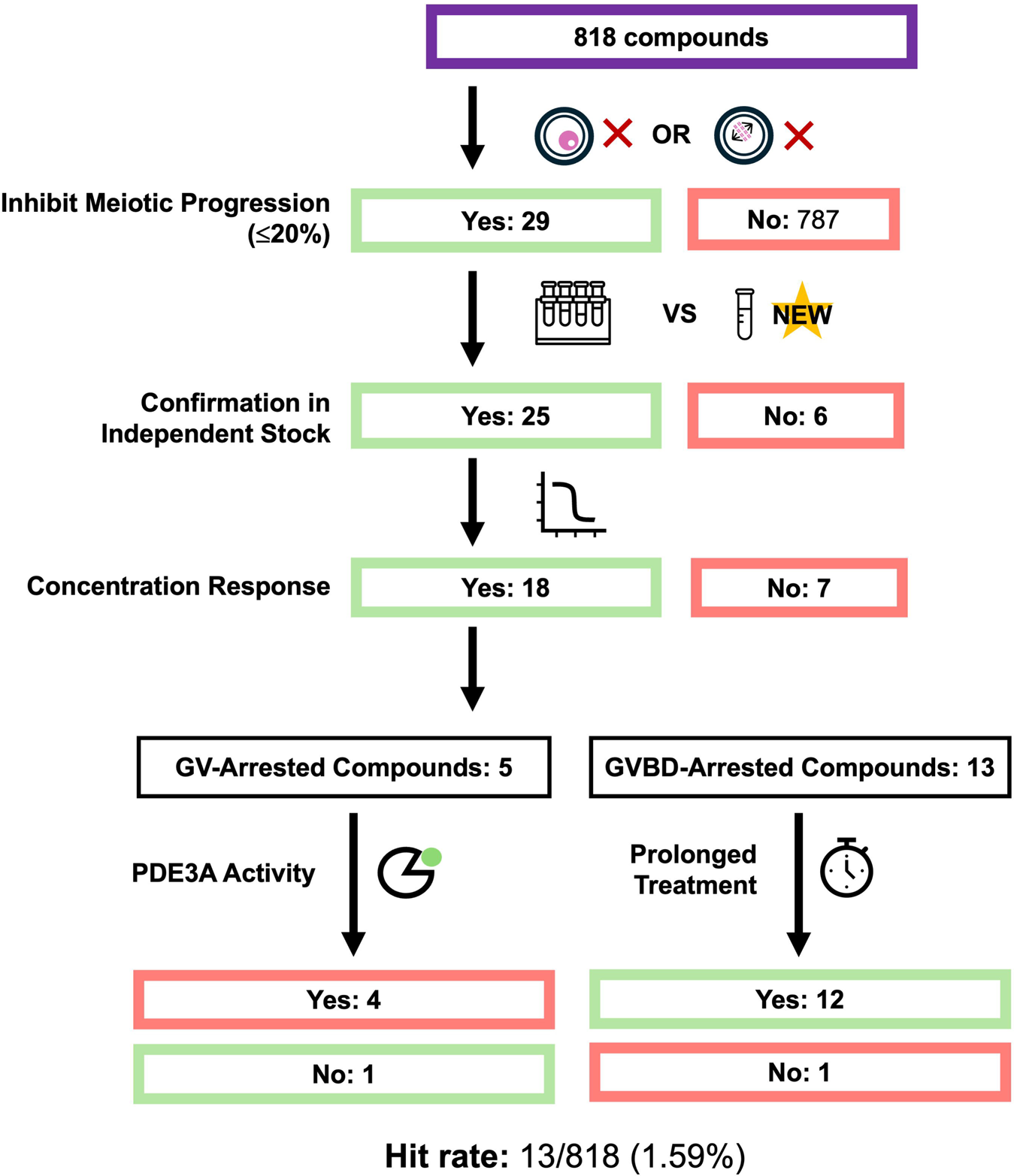
Summary of phenotypic oocyte maturation screening assay and pipeline. Hit compounds identified from the initial counterscreen assay underwent several hit validation steps, including confirmation using an independent source, concentration-response, and prolonged culture treatment. For GV-arresting compounds, an additional PDE3A activity counterscreen assay was conducted. In total, 13 compounds (1 GV-arrested and 12 GVBD-arrested) were identified through the screening pipeline. GV, germinal vesicle; GVBD, germinal vesicle breakdown; MII, meiosis II.

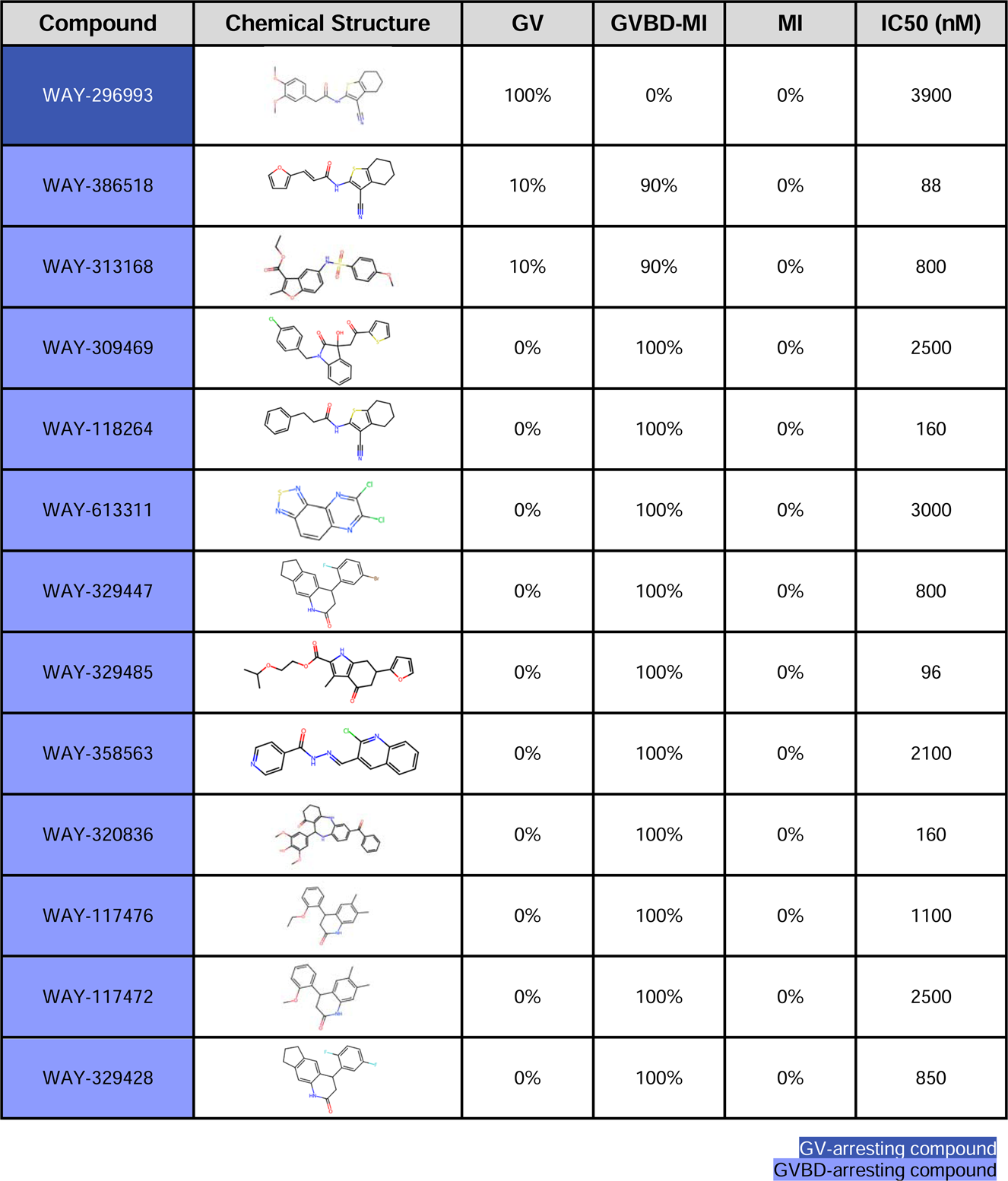

| Compound | Chemical Structure | GV | GVBD-MI | MI | IC50 (nM) |
| --- | --- | --- | --- | --- | --- |
| WAY-296993 |  | 100% | 0% | 0% | 3900 |
| WAY-386518 |  | 10% | 90% | 0% | 88 |
| WAY-313168 |  | 10% | 90% | 0% | 800 |
| WAY-309469 |  | 0% | 100% | 0% | 2500 |
| WAY-118264 |  | 0% | 100% | 0% | 160 |
| WAY-613311 |  | 0% | 100% | 0% | 3000 |
| WAY-329447 |  | 0% | 100% | 0% | 800 |
| WAY-329485 |  | 0% | 100% | 0% | 96 |
| WAY-358563 |  | 0% | 100% | 0% | 2100 |
| WAY-320836 |  | 0% | 100% | 0% | 160 |
| WAY-117476 |  | 0% | 100% | 0% | 1100 |
| WAY-117472 |  | 0% | 100% | 0% | 2500 |
| WAY-329428 |  | 0% | 100% | 0% | 850 |

GV-arresting compound GVBD-arresting compound

### Preliminary structure-activity relationships in oocyte meiotic maturation assay

During the drug discovery process, analogs of hit compounds are typically tested to determine whether specific structural features are associated with the biological activity of the hit. These structure-activity relationship (SAR) studies guide towards novel compounds whose structures have improved biological and chemical properties, such as improved potency and bioavailability as well as reduced toxicity (Guha, 2013). Furthermore, exploration of SARs allows for the identification of additional hits with known targets and thus understanding of the potential mechanism-of-action. In our curation of the 818 compounds selected for the oocyte maturation screening assay, we included structural analogs or compounds with similar chemical structures (**Supplemental Fig. 1**). Thus, we were able to compare the biological response to compounds that were identified as hits with that elicited by structurally similar compounds. Through our screening strategy, we identified a series of compounds that shared a chemical substructure but exhibited different responses in our oocyte maturation assay (**Fig. 8**). Notably, we identified one compound that resulted in GV arrest (WAY-296993) and two structurally similar compounds that resulted in GVBD arrest (WAY-118264, WAY-356518) (**Fig. 8**). In fact, WAY-296933 was originally tested as a structural analog for WAY-118264. Notably, other structural analogs (WAY-297023, WAY-614925, WAY-234384, WAY-6324922) with an additional phenyl ring did not block meiotic progression (**Fig. 8**). This preliminary SAR revealed that the compound that arrested oocytes at the GV stage (WAY-296993) had a target annotation of heat shock protein 90 (HSP90) (Chan *et al*., 2012). Our phenotype of meiotic arrest was consistent with previous work indicating that inhibition of HSP90 delays meiotic maturation in mouse and porcine oocytes (Metchat *et al*., 2009; Son *et al*., 2010; Liu *et al*., 2018). However, when we tested a series of HSP90-specific inhibitors, we did not recapitulate the observed GV arrest with WAY-296993 (**Supplemental Fig. 3**), suggesting that the compound is likely not targeting HSP90. Overall, these findings indicate the ability for our oocyte compound screening assay to develop a framework for establishing preliminary SARs and allow for ranking of biological potency based upon specific chemical structures. This approach thus allows for novel compound synthesis to generate compounds with chemical substructures that more potently block meiotic progression.

**Figure 8.**
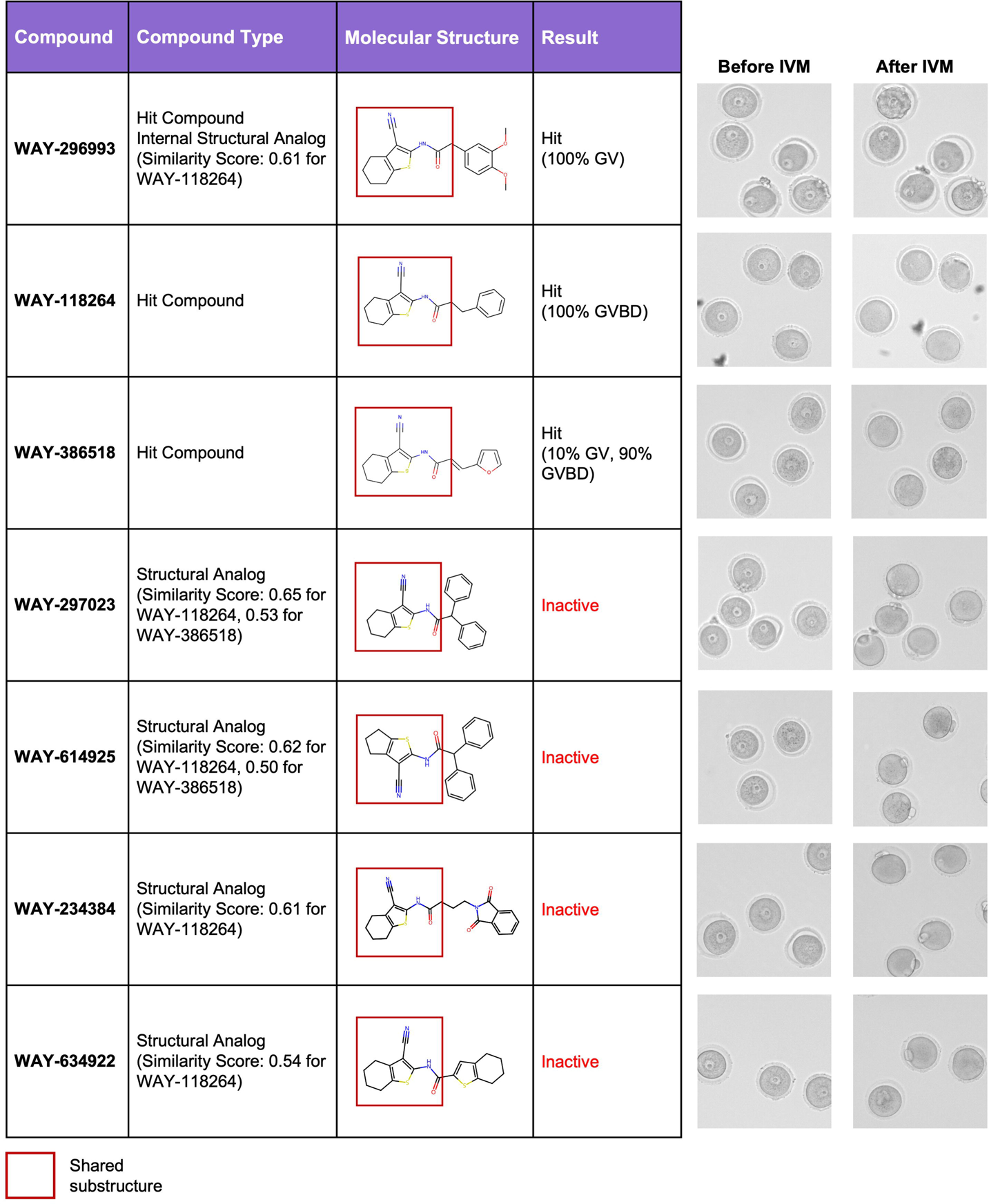
Preliminary structure-activity relationship within the phenotypic oocyte maturation assay. Table of oocyte maturation status for series of tested compounds sharing the same chemical substructure (red box). Representative images of oocytes treated with compound series before and after in vitro maturation. GV, germinal vesicle; GVBD, germinal vesicle breakdown; MII, meiosis II.

## Discussion

In this study, we established a robust oocyte maturation phenotypic screening platform using biopharmaceutical industry practices to identify compounds that block meiotic progression for female non-hormonal contraceptive development. Phenotypic drug discovery approaches offer several advantages, including the ability to identify active molecules in a target-agnostic manner and the identification of previously unknown targets (Moffat *et al*., 2017). Although high-throughput phenotypic screening systems have been previously developed in reproductive biology, they have primarily focused on targeting uterine contractility (Herington *et al*., 2015), endometriosis (Churchill *et al*., 2023), and sperm motility and function for male contraceptive development (Martins da Silva *et al*., 2017; Gruber *et al*., 2020, 2023). Given that oocytes are arrested at the GV stage throughout the reproductive lifespan and only resume meiosis during the narrow window of ovulation (Kratka *et al*., 2025), maintenance of meiotic arrest offers a promising mechanism to target for female non-hormonal contraception. Furthermore, these changes in oocyte maturation can be captured and assessed *ex vivo*, thus allowing for the opportunity to develop an assay to rapidly screen chemicals for the desired phenotype. Although our assay has lower throughput compared to other cell-based or biochemical approaches given the need to collect oocytes from hyperstimulated mice, it remains relatively simple and directly interrogates relevant biological processes in the cell-of-interest. Given the recent improvements in oocyte cryopreservation techniques (Practice Committees of the American Society for Reproductive Medicine and Society of Reproductive Biologists and Technologists, 2021), there remains an opportunity to scale up the screening platform using vitrified oocytes. Additionally, the platform does not require the use of a mineral oil overlay which is conventionally used for *in vitro* fertilization or embryo culture and can extract small molecule compounds and affect their effective concentrations (Rémillard-Labrosse *et al*., 2024).

During the development of our oocyte screening pipeline, we adopted conventional practices from the pharmaceutical industry to ensure a robust and high-quality assay. Several validation and counterscreen measures were incorporated to prevent detection of non-selective hits. Using a select number of PAINS and cytotoxic compounds, we showed that these chemicals can induce degeneration and altered cell morphology. As such, we filtered out PAINS compounds and those with unprogressable chemical features from the bioactive compound library. For hit compounds that elicited a GV-arrest phenotype, an additional counterscreen assay was conducted to determine PDE3A activity. PDE3A acts as a canonical regulator of meiotic resumption through its degradation of high cAMP levels at the GV stage (Richard *et al*., 2001; Shitsukawa *et al*., 2001). To identify novel pathways involved in regulating meiotic maturation, compounds that demonstrated PDE3A activity were deprioritized from further validation. Notably, PDE3A itself remains a promising non-hormonal contraceptive target that has been extensively evaluated across several mammalian models. Mice lacking Pde3a exhibit female infertility (Masciarelli *et al*., 2004b), and rodents treated with PDE3 inhibitors exhibited a reversible contraceptive effect without affecting ovulation or estrous cyclicity (Wiersma *et al*., 1998; Li *et al*., 2012). *In vitro* studies also confirm that pharmacological inhibition of PDE3 blocks meiotic progression in porcine and human oocytes (Nogueira *et al*., 2003; Sasseville *et al*., 2006). Lastly, treatment with a specific PDE3 inhibitor, ORG9935, has demonstrated efficacy in maintaining meiotic arrest in eggs collected from ovarian hyperstimulated rhesus macaques (Jensen *et al*., 2005, 2008). Although ORG9935 was not advanced further due to limited bioavailability and non-specific side effects during a prolonged breeding trial in macaques (Jensen *et al*., 2010), PDE3A remains a feasible target for female non-hormonal contraception, with the primary safety concern being potential impacts on increased heart rate upon inhibition (Wiersma *et al*., 1998; Jensen *et al*., 2010). Indeed, the four compounds we identified through our screening pipeline that maintained meiotic arrest through PDE3A activity represent new chemical matter that may enable development of more potent and selective PDE3A inhibitors.

Through our oocyte compound screening pipeline, we identified potent compounds that blocked meiotic progression at both the GV and GVBD stage. For oocytes arrested at the GVBD stage, most displayed scattered chromosomes and the absence of a meiotic spindle suggesting disruptions in the oocyte cytoskeleton. Notably, these phenotypes were observed despite very different compound structures (**Table 1**), suggesting that arrests at this stage typically result from disruptions in the meiotic spindle. Although it remains unclear which aspects of meiotic spindle assembly are disrupted by these compounds, they may be involved in either chromosome-mediated or microtubule-organizing center (MTOC)-mediated processes (Namgoong and Kim, 2018). Meiotic arrest due to disruptions in spindle assembly would likely lead to inability for fertilization or embryo development pre-implantation. However, targeting oocyte cytoskeletal proteins would be undesirable for non-hormonal contraception due to significant safety risks, as the chromosomal aberrations and cytoskeletal abnormalities would also likley impact somatic cells resulting in high risk for carcinogenesis. Furthermore, unlike GV oocytes, GVBD oocytes can still be fertilized and generate triploid embryos which consequently leads to spontaneous abortions (Lauritsen *et al*., 1979; McFadden and Langlois, 2000). As such, future efforts in scaling up compound screening in oocytes would likely deprioritize compounds that elicit arrest at the GVBD stage. Rather, these compounds may be leveraged to identify new mechanisms in oocyte biology associated with meiosis and fertility.

Although we were able to identify potent compounds that blocked meiotic progression, we did not identify which specific targets that may underlie the observed phenotype. Indeed, despite one of our GV-arresting compounds (WAY-296993) being annotated to target HSP90 in somatic cells, we could not recapitulate the phenotype using HSP90-specific inhibitors. These results indicate that annotated targets based on literature require comprehensive validation to confirm their role in oocytes. Future studies are thus required to conduct target deconvolution, which can be determined through chemical, genomic, and proteomic approaches. Given that our oocyte screening assay can detect phenotypic differences across structurally similar compounds, testing of additional structural analogs would allow for development of an expanded SAR. Transcriptomic profiling of a given cell type or biological system in response to many chemical perturbations allow for the creation of a perturbome. Indeed, evaluation of perturbation gene signatures between known tool compounds and unknown compounds allows for identification of drug mechanism-of-action and new drug targets (Caldera *et al*., 2019; Szalai and Veres, 2023). Proteomic approaches of target deconvolution, such as affinity chromatography or phage display, commonly utilize immobilized compounds to allow for binding of partner proteins and their subsequent detection using mass spectrometry or DNA sequencing, respectively (Terstappen *et al*., 2007). More recently, thermal proteomic profiling has been developed which leverages the improved stability of small molecule-bound proteins and compares the melting point of proteins with and without treatment for target identification (Franken *et al*., 2015; Mateus *et al*., 2020).

In summary, we developed a phenotypic screening platform for oocyte meiotic maturation and established a robust pipeline for screening compounds using conventional industry drug discovery practices. Through this process, we identified potent compounds that resulted in precise biological phenotypes in blocked meiotic progression. This study also demonstrated that the inhibitory effect of these compounds can persist throughout the window of fertilization and be a reversible process. Therefore, we highlight this methodology and the identified hit compounds as a starting point for target deconvolution and development of drug discovery programs for female non-hormonal contraception.

## Author’s Roles

J.P., Y.Z., H.C.L., and F.E.D. designed the experiments. Y.Z., H.C.L., and L.R.H. performed the experiments. Y.Z., L.R.H., and J.P. analyzed the data. All authors provided critical feedback and contributed to the interpretation of the results. J.P. prepared the original draft of the manuscript and all authors reviewed and edited the manuscript. All authors have read and agreed to the published version of the manuscript.

## Acknowledgements

We thank the members of the Duncan lab and the Ovarian Contraceptive Discovery Initiative for their support and advice regarding this work.

## Funding

This work was supported by the Gates Foundation [INV-003385]. Under the grant conditions of the Foundation, a Creative Commons Attribution 4.0 Generic License has already been assigned to the Author Accepted Manuscript version that might arise from this submission.

## Conflicts of interest

The authors have no conflict of interest to disclose.

## Notes

### Competing Interest Statement

The authors have declared no competing interest.

