## Supplemental Figures for "A complex phenotypic assay of mammalian oocyte maturation identifies compounds that block meiotic progression for non-hormonal contraceptive discovery"

### Slide 1
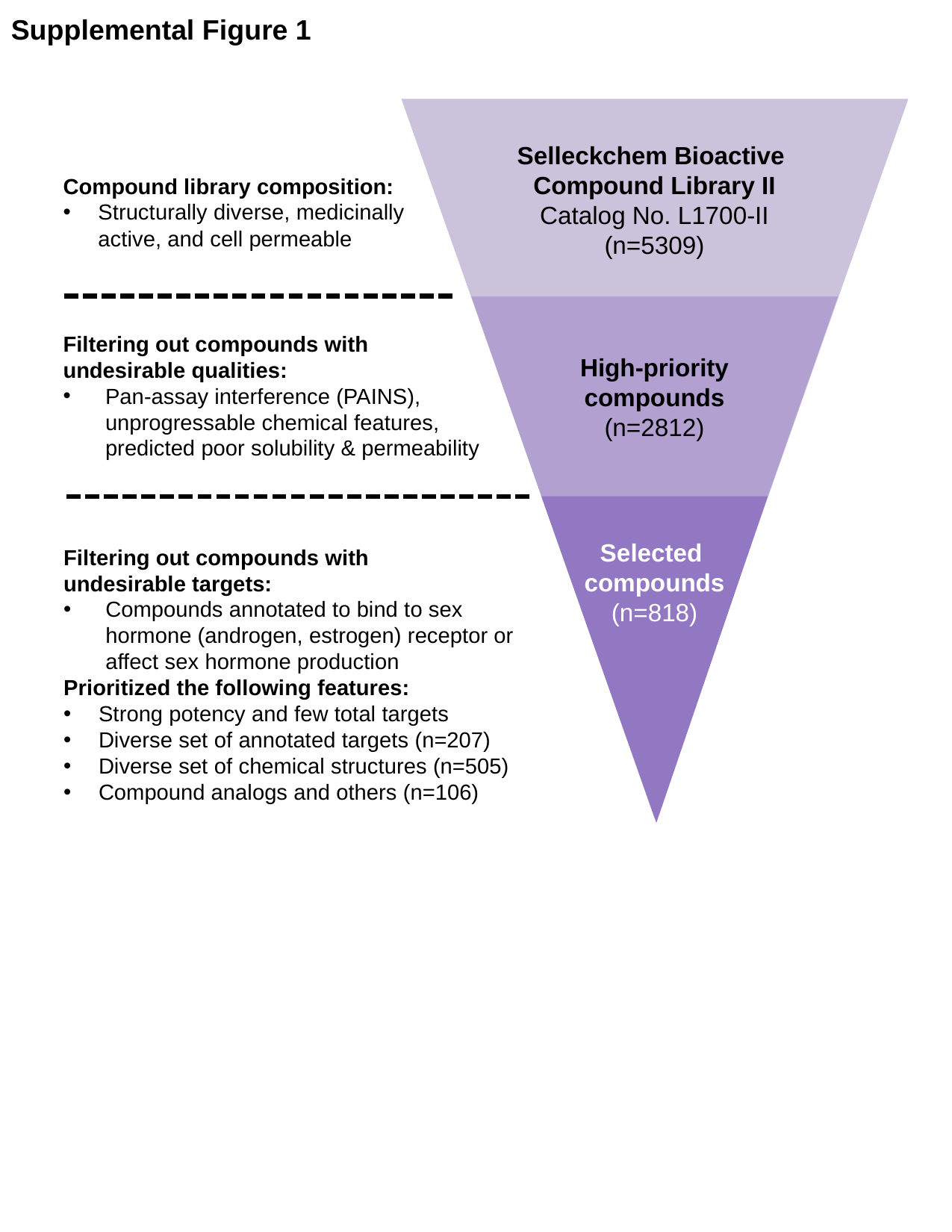

Supplemental Figure 1
Selleckchem Bioactive
Compound Library II
Catalog No. L1700-II
(n=5309)
High-priority
compounds
(n=2812)
Selected
compounds
(n=818)
Compound library composition:
Structurally diverse, medicinally active, and cell permeable
Filtering out compounds with undesirable qualities:
Pan-assay interference (PAINS), unprogressable chemical features, predicted poor solubility & permeability
Filtering out compounds with
undesirable targets:
Compounds annotated to bind to sex hormone (androgen, estrogen) receptor or affect sex hormone production
Prioritized the following features:
Strong potency and few total targets
Diverse set of annotated targets (n=207)
Diverse set of chemical structures (n=505)
Compound analogs and others (n=106)

### Slide 2
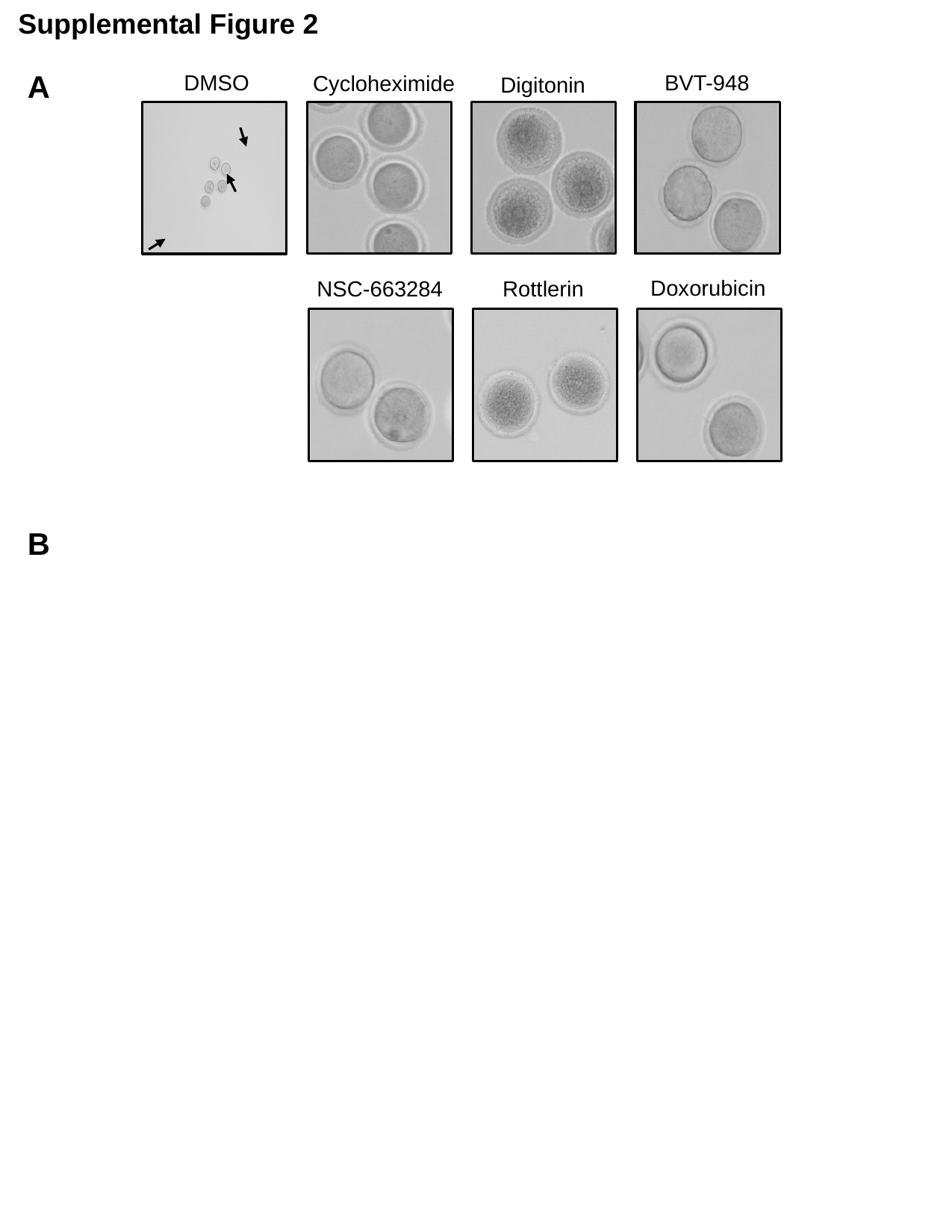

Supplemental Figure 2
A
DMSO
BVT-948
Cycloheximide
Digitonin
Doxorubicin
Rottlerin
NSC-663284
B

### Slide 3
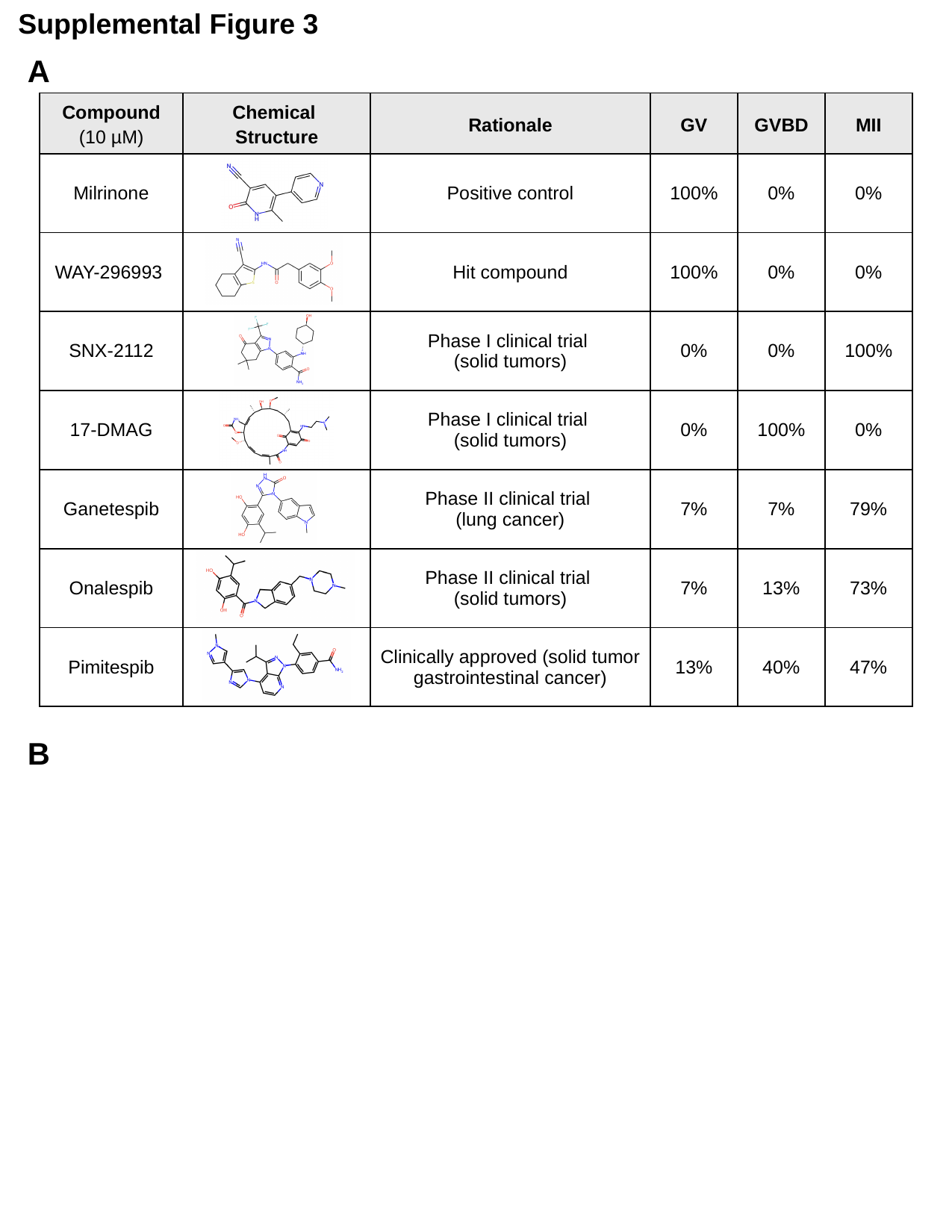

Supplemental Figure 3
A
| Compound (10 µM) | Chemical Structure | Rationale | GV | GVBD | MII |
| --- | --- | --- | --- | --- | --- |
| Milrinone​ | | Positive control | 100%​ | 0%​ | 0%​ |
| WAY-296993 ​ | | Hit compound | 100%​ | 0%​ | 0%​ |
| SNX-2112 | | Phase I clinical trial (solid tumors) | 0%​ | 0%​ | 100% |
| 17-DMAG | | Phase I clinical trial (solid tumors) | 0%​ | 100%​ | 0%​ |
| Ganetespib | | Phase II clinical trial (lung cancer) | 7%​ | 7%​ | 79%​ |
| Onalespib | | Phase II clinical trial (solid tumors) | 7% | 13% | 73% |
| Pimitespib | | Clinically approved (solid tumor gastrointestinal cancer) | 13% | 40% | 47% |
B
